# PLK1 plays dual roles in centralspindlin regulation during cytokinesis

**DOI:** 10.1101/317388

**Authors:** Ingrid E. Adriaans, Angika Basant, Bas Ponsioen, Michael Glotzer, Susanne M. A. Lens

## Abstract

Cytokinesis starts in anaphase with the formation of an actomyosin-based contractile ring at the equatorial cortex, which is governed by the local activation of the small GTPase RhoA. Here we delineated the contributions of PLK1 and Aurora B to RhoA activation and cytokinesis initiation in human cells. Knock-down of PRC1, which disrupts the spindle midzone, revealed the existence of two pathways that can initiate cleavage furrow ingression. One pathway depends on a well-organized spindle midzone and PLK1, while the other depends on Aurora B activity and centralspindlin oligomerization at the equatorial cortex and can operate independently of PLK1. We further show that PLK1 inhibition sequesters centralspindlin onto the spindle midzone making it unavailable for Aurora B-dependent oligomerization at the equatorial cortex. We propose that PLK1 activity promotes the release of centralspindlin from the spindle midzone through inhibition of PRC1, allowing centralspindlin to function as a regulator of spindle midzone formation and as an activator of RhoA at the equatorial cortex.

## Introduction

Cytokinesis drives the physical separation of daughter cells at the end of mitosis. Failure to complete cytokinesis gives rise to tetraploid cells with supernumerary centrosomes. Depending on the cell type and cellular context, cytokinesis failure can either result in a G1 arrest, or allow cell cycle progression of the tetraploid cells into the next mitosis (Uetake and Sluder, 2004; Andreassen et al., 2001). These dividing tetraploid cells are at risk of becoming aneuploid, owing to, for example, the extra number of centrosomes that can cause the missegregation of chromosomes during mitosis (Ganem et al., 2007; Tanaka et al., 2015; Silkworth et al., 2009). Hence proper execution and completion of cytokinesis is essential for genomic stability.

In animal cells, cytokinesis starts in anaphase with the formation of an actomyosin-based contractile ring at the equatorial cortex that drives ingression of the cleavage furrow. Prior to membrane furrowing interpolar microtubules are bundled between the separating sister chromatids to form the spindle midzone (also referred to as central spindle). As the furrow ingresses, these microtubule bundles are compacted into a cytoplasmic bridge, with the midbody in its center. The midbody attaches the ingressed cell membrane to the intercellular bridge and promotes the final phase of cytokinesis, known as abscission (Lekomtsev et al., 2012; D’Avino and Capalbo, 2016; Steigemann and Gerlich, 2009; Hu et al., 2012b). Formation of the contractile ring requires activation of the small GTPase RhoA by the Guanine Nucleotide Exchange Factor (GEF), ECT-2 (Basant and Glotzer, 2018). Active, GTP-bound RhoA assembles and activates components of the actomyosin-based ring, such as diaphanous-related formin that facilitates the assembly of actin filaments (Otomo et al., 2005; Watanabe et al., 2008; Piekny et al., 2005; Chen et al., 2017), and Rho-Kinase (ROCK), which activates nonmuscle myosin II to power ring constriction (Kosako et al., 2000; Amano et al., 1996). Optogenetic manipulation of RhoA activity showed that local activation of RhoA on the cell membrane is sufficient to drive cleavage furrow initiation independent of cell cycle stage (Wagner and Glotzer, 2016). Hence, strict spatial and temporal regulation of RhoA activity is essential to coordinate the onset of cytokinesis with nuclear division.

Current models for local RhoA activation and cleavage furrow initiation involve at least two anaphase spindle-derived stimulatory signals: one originating from the spindle midzone and another derived from astral microtubules that end at the equatorial cortex (Mishima, 2016). Experiments in large echinoderm embryos suggest a stimulatory role of astral microtubules in the initiation of cleavage furrow ingression (Mishima, 2016; Su et al., 2014), whilst data in smaller (mostly mammalian) cells emphasized a role for the spindle midzone (Cao and Wang, 1996). The overlapping antiparallel microtubules of the spindle midzone serve as a platform for the localization of a variety of proteins that promote RhoA activation and cleavage furrow ingression directly parallel to the microtubule overlap. In addition, astral microtubules convey inhibitory signals at cell poles (Mangal et al., 2018; Werner et al., 2007; Wagner and Glotzer, 2016; Su et al., 2011; Kotýnková et al., 2016).

Protein Regulator of Cytokinesis 1 (PRC1) is essential for the assembly of a fully functional spindle midzone (Zhu et al., 2006; Mollinari et al., 2005, 2002). PRC1 is a homodimeric microtubule bundling protein that as a homodimer binds the interface between antiparallel microtubules (Li et al., 2018). Its microtubule bundling activity is required for spindle midzone formation, thereby indirectly contributing to the recruitment of other spindle midzone-localized proteins, such as centralspindlin and the Chromosomal Passenger Complex (CPC) (Zhu et al., 2006; Mollinari et al., 2005). Furthermore, through interaction with the kinesin KIF4A and Polo-like kinase 1 (PLK1) (Zhu and Jiang, 2004; Kurasawa et al., 2004), PRC1 also acts as a direct recruiter of regulatory proteins to the spindle midzone.

Centralspindlin is a heterotetramer consisting of two molecules of the kinesin-6 MKLP1 (KIF23) and two molecules of RACGAP1 (hsCyk4, MgcRacGAP) (Basant and Glotzer, 2018). Oligomerization of the complex is needed to bundle microtubules and organize the spindle midzone (Hutterer et al., 2009). In addition to microtubule bundling, centralspindlin also promotes RhoA activation and cleavage furrow initiation (Somers and Saint, 2003; Yüce et al., 2005;Nishimura, 2006). This latter function of centralspindlin appears to rely on PLK1-dependent binding of RACGAP1 to ECT-2 (Petronczki et al., 2007; Burkard et al., 2009; Wolfe et al., 2009). Mutation of PLK1 phosphorylation sites in the non-catalytic N-terminus of RACGAP1 disrupts its ability to interact with the N-terminal BRCT domain in ECT-2 and disturbs the initiation of cytokinesis (Burkard et al., 2009; Wolfe et al., 2009). In line, inhibition of PLK1 activity at anaphase onset prevents RhoA activation at the equatorial cortex and cleavage furrow ingression (Petronczki et al., 2007; Brennan et al., 2007). Because inhibition of PLK1 activity or expression of a RACGAP1 PLK1-phosphorylation site mutant disrupt the spindle midzone localization of ECT-2, it was proposed that the spindle midzone localization of ECT-2 is a determining factor for the spatial activation of RhoA at the equatorial cortex (Burkard et al., 2009; Wolfe et al., 2009).

The Chromosomal Passenger Complex (CPC), consisting of INCENP, Survivin, Borealin and Aurora B kinase, relocates from chromosomes to the spindle midzone and equatorial cortex at anaphase onset in an MKLP2 (KIF20A)-dependent manner (Gruneberg et al., 2004; Hümmer and Mayer, 2009; Kitagawa et al., 2013, 2014). One of the spindle midzone targets of Aurora B is the centralspindlin subunit MKLP1 (Guse et al., 2005; Neef et al., 2006). Phosphorylation of S708 of MKLP1 disrupts the interaction between MKLP1 and 14-3-3 proteins, resulting in oligomerization of centralspindlin (Guse et al., 2005; Douglas et al., 2010; Basant et al., 2015). In *C. elegans* embryos, Aurora B-induced oligomerization of centralspindlin promotes RhoA activation and cleavage furrow ingression and Aurora B activity is largely dispensable when centralspindlin constitutively oligomerizes (Basant et al., 2015). However, in mammalian cells, inhibition of Aurora B kinase activity at anaphase onset does not prevent initiation of cytokinesis, though it does prevent completion of cytokinesis (Guse et al., 2005; Ahonen et al., 2009). This suggests that RhoA activation can occur in the absence of Aurora B activity in mammalian cells, and that in these cells Aurora B activity appears to be more relevant at later stages of cytokinesis most likely by promoting the formation of the spindle midzone and midbody (D’Avino and Capalbo, 2016).

Whilst the spindle midzone is considered to provide important cues for RhoA activation at the equatorial cortex in mammalian cells (Yüce et al., 2005;Nishimura, 2006; Petronczki et al., 2007; Burkard et al., 2009; Wolfe et al., 2009), knock-down of PRC1, which clearly disrupts the spindle midzone, does not impair RhoA activation and cleavage furrow ingression (Jiang et al., 1998;Mollinari et al., 2005; Zhu and Jiang, 2004; Zhu et al., 2006). This implies that spindle midzone-independent cues can also specify and activate RhoA in small, mammalian cells. Here we demonstrate that in the absence of PRC1, RhoA is activated at the equatorial cortex through centralspindlin oligomerization induced by cortical Aurora B activity. Remarkably, we find that in PRC1-deficient cells, cytokinesis initiation can occur in the absence of PLK1 activity. This alternative PLK1-independent route to RhoA activation has been overlooked because of an unrecognized inhibitory effect of PLK1 on PRC1 in anaphase. We find that PLK1 activity limits PRC1-dependent hyper-bundling of spindle midzone microtubules, and reduces centralspindlin sequestration at the spindle midzone, making it available to activate ECT-2 at the equatorial cortex.

## Materials and Methods

### Cell culture

HeLa cells and HeLa Flp-In T-Rex cells were cultured in DMEM (Sigma-Aldrich) supplemented with 6% FCS (FBS, Sigma-Aldrich), 2 mM UltraGlutamine and 100 units/ml penicillin and 100 μg/ml streptomycin (Lonza). Culture medium of HeLa Flp-In T-Rex cells and of all HeLa Flp-In T-Rex derived cell lines (described below) was additionally supplemented with 4μg/ml Blasticidin (PAA Laboratories). All cell lines were cultured at 37°C with 5% CO_2_.

### Plasmids

cDNA encoding the membrane-binding Tubby domain of mouse Tubby protein (aa 243-505) (Szentpetery et al., 2009) was obtained by PCR from a mouse brain cDNA library and was subsequently cloned into a pcDNA3 vector containing a mTFP1 and YPet FRET based sensor for Aurora B activity (Fuller et al., 2008; Wurzenberger et al., 2012). The CyPet donor of the original construct was replaced by mTFP1. Full length, wild-type MKLP1 was amplified by PCR from a human thymus cDNA library and cloned into a pEGFP-C1 vector (Clontech) and subsequently cloned into a pcDNA5^TM^/FRT/TO vector (Invitrogen) in which the hygromycin B resistance cassette was replac by a puromycin B resistance cassette (pcDNA5/FRT/TO-puro). Point mutations were introduced by site-directed mutagenesis.

### siRNA and plasmid transfection

HeLa cells were transfected with siLUC (Luciferase GL2 duplex;Dharmacon/D-001100- 01-20), siPRC1 (ON-TARGETplus SMARTpool; L-019491-00-0005) siMKLP2 (ON-TARGETplus SMARTpool; L-019491-00-0005), siMKLP1 (CGACAUAACUUACGACAAAUU) or siRACGAP1 (Thermo Fischer, HSS120934 Stealth siRNA) with HiPerfect Transfection Reagent (#301705; Qiagen). The final concentration of siRNAs was 20nM for siLUC, siPRC1, RACGAP1 and MKLP2. The final concentration of siRNAs for siMKLP1 was 40nM. Cells were analyzed 48 hours after siRNA transfection. A standard HiPerfect transfection protocol was used with a 3:1 ratio for siRNA:HiPerfect in Opti-MEM. Incubation of siRNA/HiPerfect mixture was done at 37°C for 20 minutes. Transient transfection of plasmids was performed with X-tremeGENE^™^ 9 DNA Transfection Reagent according the manufacturers protocol (Roche). To generate stable cell lines with doxycyclin-inducible expression of GFP::MKLP1 and GFP::MKLP1-S708E, HeLa Flp-In T-Rex cells were co-transfected with pOG44 (Invitrogen) and pcDNA5/FRT/TO-puromycin plasmids encoding the indicated MKLP1 proteins. After transfection cells were selected in medium supplemented with 2μg/ml puromycin and 4μg/ml blasticidin (Invitrogen). Finally, to generate HeLa Flp-In T-Rex cells stably expressing the Tubby-Aurora B FRET sensor, cells were transfected with a pcDNA3 plasmid encoding Tubby-Aurora B FRET sensor and cells were selected and maintained in medium supplemented with 2μg/ml puromycin (Sigma-Aldich) and 4μg/ml blasticidin (Invitrogen).

### Cell synchronization and inhibitor treatment

For live cell imaging and immunofluorescence of anaphase cells, HeLa cells were plated in 2.5mM thymidine (Sigma-Aldrich, cat. no. T1895) for 24 hours and released into 5μM RO3306 (Calbiochem) for 16 hours. Where indicated, doxycycline (1 μg/m, Sigma-Aldrich) was added together with RO3306 (Calbiochem). Cells were released from the RO3306 block by washing three times with medium. HeLa cells were either filmed immediately after release from RO3306 or fixed 60 minutes after RO3306 release. Where indicated BI2536 (100 nM final concentration, Selleck Chemicals) or ZM447439 (2 μM final concentration, Tocris Bioscience) was added 35 minutes after the RO3306 release. Alternatively, 83 nM DMSO or nocodazole (Sigma-Aldrich) was added 45 minutes after RO3306 release.

### Western blot sample preparation, SDS-PAGE and Western blotting

HeLa cells were synchronized in G1/S phase by a 24 h incubation with thymidine. After release from the thymidine block, cells were accumulated in mitosis by addition of S-Trityl-L-Cysteine (STLC, 20 μM, Tocris Bioscience) for 16 hours. Mitotic enriched cells were collected and lysed in Laemmli buffer. Protein concentration was determined using a Lowry assay and protein samples were separated by SDS-PAGE and transferred to nitrocellulose membranes. Membranes were blocked in 4% milk in Tris-buffered Saline containing 0.5% Tween-20 (TBST) and subsequently incubated with the primary antibody for 2 hrs. Primary antibodies used were rabbit anti-PRC1 (sc-8356), mouse anti-alpha tubulin (T5168), rat anti-RFP (Chromotek 5F8), goat anti-RACGAP1 (Abcam ab2270), rabbit anti-MKLP1 (sc-867) and mouse anti-GFP (Roche). After washing the membranes with TBST, they were incubated with horseradish peroxidase (HRP)-conjugated secondary antibodies (Bio-Rad). An ECL chemiluminescence detection kit (GE Healthcare) was used to visualize the protein/antibody complex.

### Immunofluorescence Microscopy

Cells were grown in 24 well plates containing 12 mm High Precision coverslips (Superior-Marienfeld GmbH & Co). Cells were fixed with 4%PFA in PBS for 7 minutes and permeabilized in either methanol (−20°C) or in 0.25% Triton X-100 in PBS for 5 minutes (RACGAP1). Cells were blocked in PBS containing 3% BSA and 0.1% Tween-20. Primary antibodies used were rabbit KIF20A (Bethyl (ITK) A300-878A), rabbit PRC1 (Santa Cruz, sc-8356), mouse Aurora B (BD Transduction labs, 611083), rabbit Anillin (Piekny & Glotzer 2008) mouse RhoA (Santa Cruz, sc-418), rabbit MKLP1 (Santa Cruz, sc-867), rabbit PLK1 (Santa Cruz, sc-17783), mouse *α*-tubulin (Thermo Fisher, MA1-80017), goat RACGAP1 (Abcam, ab2270). Secondary antibodies used were goat anti-mouse or goat anti-rabbit IgG-Alexa 488, goat anti-mouse or goat anti-rabbit IgG-Alexa 568 and goat anti-rat Alexa 647 (Invitrogen).

In case of RACGAP1 staining, donkey anti-goat IgG-Alexa 568 was used in combination with chicken anti-mouse IgG-Alexa 488. For IF of RhoA, cells were fixed with 10% Trichloroacetic acid (TCA) in H_2_O for 15 minutes on ice, washed 3 times with 30 mM glycine in PBS and permeabilized for 5 minutes at room temperature in 0.25% Triton-X 100 in PBS. Antibody dilutions and washes were done in 5% non-fat milk in PBS. 4’,6-Diamidino-2-Phenylindole (DAPI) was used for DNA staining and coverslips were mounted using ProLong Antifade (Molecular Probes). Images were taken with a Personal DeltaVision system (Applied Precision) equipped with a 40 × /1.35 numerical aperture (NA) UPlanSApo objective (Olympus) and a CoolSNAP HQ camera (Photometrics).

### Live cell microscopy

Cells were seeded in a μ-Slide 4 well (ibiTreat, Ibidi) and medium was changed to Leibovitz’s medium (Sigma) supplemented with 10% FCS (FBS, Sigma-Aldrich), 2 mM UltraGlutamine and 100 units/ml penicillin and 100 μg/ml streptomycin (Lonza) prior to live cell imaging. FRET sensor imaging was performed on an inverted NIKON Diaphot microscope (Nikon-Eclipse Ti) with a Perfect Focus System (PFS). Imaging was done with a 63x OIL N.A. 1.49 WD 0.12 objective. Imaging was performed at 37°C using a Microscope Cage Incubator (OkoLab). The excitation filter, dichroic mirror and emission filter for mTFP1 were 430/24x, Z465LP and 480/40m. The excitation filter, dichroic mirror and emission filter for YFP were 500/20x, 89006bs and 535/30m (Chroma). The emissions from both fluorophores were collected simultaneously from a single Z plane using a TuCam beamsplitter device (Andor). Other fluorescent live cell imaging experiments were performed with a Personal DeltaVision system (Applied Precision) with a 63 × /1.35 numerical aperture (NA) UPlanSApo objective (Olympus) using a CoolSNAP HQ camera (Photometrics). DIC images were acquired using an Olympus IX-81 microscope with a Hamamatsu ORCA-ER CCD Camera and a 37°C heated chamber, controlled by Cell-M software. Image analysis was performed with Fiji image processing software (ImageJ).

### FRET analysis

FRET sensor data analysis was done with a customized Image J FRET macro. Background emissions were measured by selecting a Region of Interest (ROI) adjacent to the cell cortex outside the cell. The background emissions were subtracted from both mTFP and Ypet emissions and the YPet/mTFP ratio was calculated. Area selection was performed using thresholding and manual selection of the equatorial cell cortex.

### C. elegans experiments

*C. elegans* strains were maintained on nematode growth medium (NGM) plates using standard procedures. RNAi was administered by feeding nematodes with E. coli expressing the appropriate double-stranded RNA (dsRNA) (Timmons and Fire, 1998). HT115 bacterial cultures were grown in Luria broth with 100 μg/ml ampicillin overnight at 37°C. Cultures (250 μl) were seeded on NGM plates containing 100 μg/ml ampicillin and 1 mM isopropyl β-d-1-thiogalactopyranoside and incubated at room temperature (~23°C) for 8 hours. RNAi plasmids were obtained from a library produced by (Kamath et al., 2003). L4 hermaphrodites were picked onto feeding plates at 25°C at least 24 hours prior to dissection. To prepare one-cell embryos for imaging, gravid hermaphrodites were dissected into egg salt buffer on coverslips, mounted onto 2.5% agar pads and sealed with vaseline. For confocal imaging, embryos were imaged with a 63X/1.4 numerical aperture oil-immersion lens on either a Zeiss Axiovert 200M equipped with a Yokogawa CSU-10 spinning-disk unit (McBain, Simi Valley, CA) and illuminated with 50-mW, 473-nm and 20-mW, 561-nm lasers (Cobolt, Solna, Sweden), or a Zeiss Axioimager M1 equipped with a Yokogawa CSU-X1 spinning-disk unit (Solamere, Salt Lake City, UT) and illuminated with 50-mW, 488-nm and 50-mW, 561-nm lasers (Coherent, Santa Clara, CA). Images were captured on a Cascade 1K EM-CCD camera or a Cascade 512BT (Photometrics, Tucson, AZ) controlled by MetaMorph (Molecular Devices, Sunnyvale, CA). Image processing was performed with ImageJ. Time-lapse acquisitions were assembled into movies using Metamorph and ImageJ. *Nop-1(it142)* embryos expressing mCherry::H2B; mCherry::PH and CYK4::GFP, or embryos expressing CYK-4::GFP and a membrane marker (mCherry::PH) were filmed simultaneously by time-lapse confocal and DIC microscopy. In the mCherry::PH or the DIC images, the tip of the furrow was defined by an ROI in ImageJ at different stages of ingression. The maximum intensity of CYK-4::GFP in the ROI was determined in the corresponding frames. Using the same ROI, a maximum intensity in a random region of the cytoplasm was obtained. The extent of recruitment to the furrow tip was calculated as (max. furrow intensity/max. cytosolic intensity −1). These values were averaged across embryos of a given genotype.

### Statistical analysis

Where indicated, the mean and standard deviation (SD) are shown. Statistical significance was calculated with a Chi-squared test or a Student’s t-test using Prism 7 software.

## Results

### Disruption of the spindle midzone allows PLK1-independent RhoA activation and furrow ingression

Cleavage furrow ingression can take place in human cells lacking PRC1 despite the absence of a well-organized spindle midzone that concentrates cytokinetic regulators (Mollinari et al., 2005; Zhu et al., 2006). This suggests the existence of spindle midzone-independent cues in human cells that contribute to the local concentration of active RhoA. To delineate this spindle midzone-independent pathway, we analyzed the contribution of key cytokinesis regulators to cytokinesis initiation in a PRC1 knock-down background. We first confirmed that after PRC1 knock-down the spindle midzone was severely disrupted, as shown by the disorganized appearance of midzone microtubules (Fig. 1A,B). In addition, live cell imaging of anaphase cells depleted of PRC1 reveals that most cells exhibit full cleavage furrow ingression followed by cleavage furrow regression (Fig. 1C,D). The frequency and extent of furrow ingression was assessed and a distinction was made between no furrow ingression and minimal furrow ingression (Fig. 1D).

**Figure 1:**
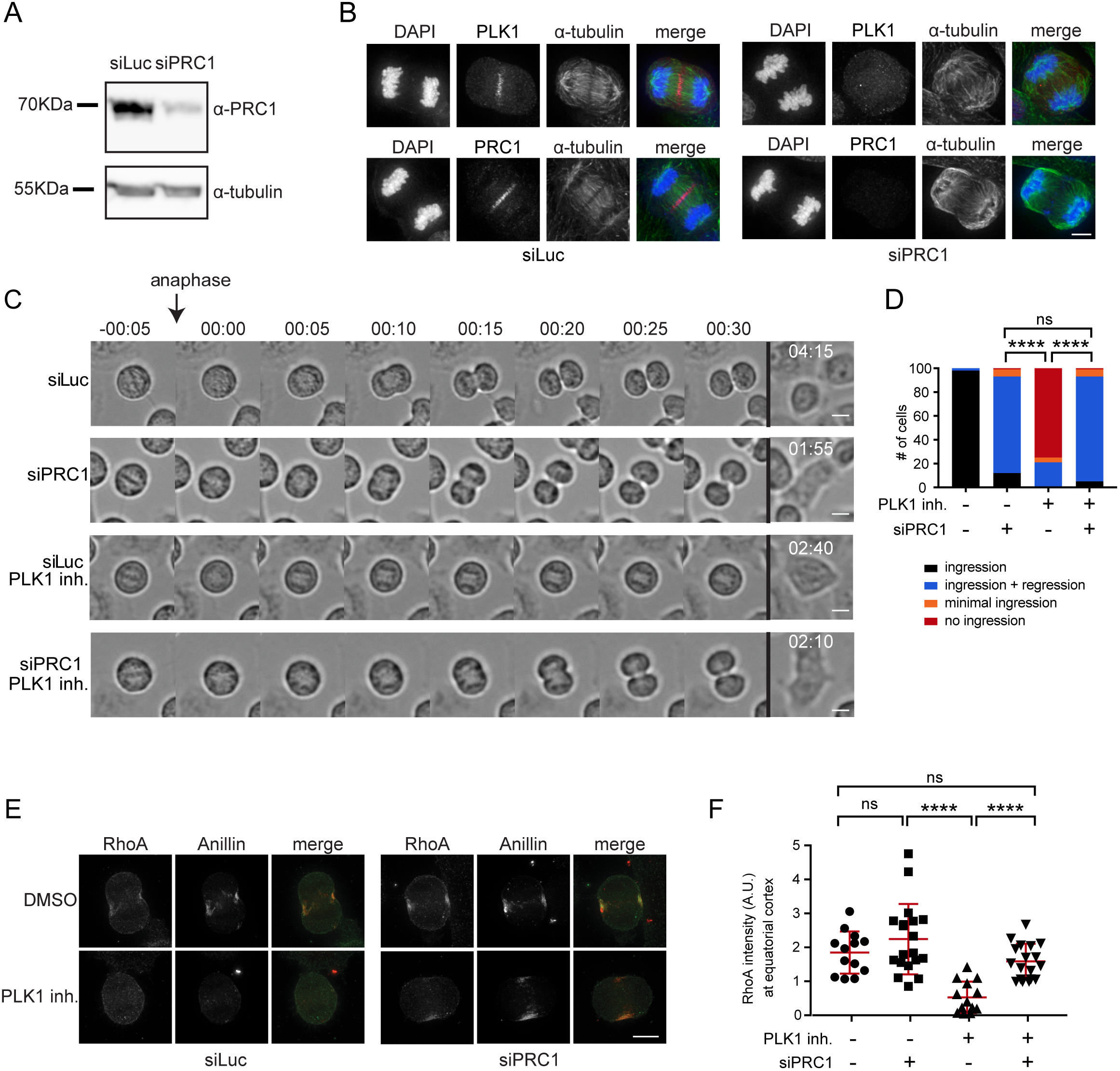
PRC1 depletion reveals PLK1-independent RhoA activation and furrow ingression. (A) Western blot of HeLa cells transfected with control siRNA (siLuc) or siRNA specific for PRC1. The Western blot was probed with an anti-PRC1 antibody. α-Tubulin is shown as loading control. (B) IF for PLK1, PRC1 and α-tubulin of HeLa cells transfected with siLuc or siPRC1. DNA was visualized using DAPI. Scale bar indicates 5 μm. (C) Representative DIC stills of a live cell imaging experiment of HeLa cells transfected with either siLuc or siPRC1 plus or minus addition of BI2536 (100 nM) at anaphase onset. Scale bar indicates 10 μm. (D) Percentage of cells showing either complete furrow regression, full furrow regression followed by regression, visible but minimal furrow ingression or no furrow ingression (n = 100 cells imaged per condition). **** = p < 0.0001; Chi-squared test for comparison of the indicated conditions, ns = not significant. One representative experiment out of 2 is shown. (E) IF for RhoA and Anillin of HeLa cells in anaphase transfected with the indicated siRNAs and treated plus or minus BI2536 (100 nM). Scale bar indicates 10 μm. (F) Quantification of fluorescence intensity levels of RhoA at the equatorial cortex in anaphase. Each dot represents an individual cell. Error bars depict the standard deviation (SD). **** = p < 0.0001; Student’s t-test for comparison of the indicated conditions, ns = not significant.

Because PLK1 is a critical regulator of RhoA activation and cleavage furrow ingression (Burkard et al., 2009; Petronczki et al., 2007), we tested its involvement in spindle midzone-independent cytokinesis initiation. PLK1 activity was inhibited at the onset of anaphase by addition of the small molecule inhibitor BI2536 (Steegmaier et al., 2007). Remarkably, while PLK1 inhibition alone completely prevented furrow ingression in ~80% of control transfected cells, PLK1 inhibition allowed full furrow ingression in PRC1 knockdown cells (Fig 1C,D), after a ± 5 min. delay in the onset of furrow ingression (Suppl. Fig. 1A). In line with this, PRC1 depletion restored RhoA and Anillin localization to the equatorial zone in PLK1 inhibited cells (Fig. 1E,F). Thus, although PLK1 activity is critical for RhoA activation and initiation of cytokinesis in PRC1 containing cells, it is dispensable when PRC1 is absent and the spindle midzone is disrupted. This suggests that PRC1 can function both as an activator and as an inhibitor of RhoA activation.

### PLK1-independent furrow ingression relies on the centralspindlin complex

Because RACGAP1 of the centralspindlin complex is an important target of PLK1 and involved in the activation of the RhoA GEF, ECT-2 (Burkard et al., 2009; Wolfe et al., 2009; Brennan et al., 2007; Santamaria et al., 2007), we next tested whether centralspindlin is required for furrow ingression in cells depleted of PRC1 and treated with an inhibitor of PLK1. After RACGAP1 knock-down the protein was indeed no longer detectable on Western blot (Fig. 2A). MKLP1 knockdown was confirmed by IF as the antibody failed to detect the protein on Western blot (Fig. 2B). The frequency of cells with undetectable MKLP1 localization in the anaphase midzone was used as a proxy for knockdown efficiency. Next, we combined PLK1 inhibition with knockdown of either PRC1, RACGAP1 or MKLP1, or with a combination of PRC1 and RACGAP1 knockdown, or PRC1 and MKLP1 knockdown (Fig. 2C-E). Approximately 80% of the siPRC1 transfected cells, initiated cytokinesis when PLK1 was inhibited (Figs 1D and 2C,D). However, cytokinesis initiation was almost completely blocked when RACGAP1 or MKLP1 was also depleted (Fig. 2C-E and Suppl. Fig. 1B). This demonstrates that whilst spindle midzone-independent furrow ingression does not require PLK1 activity, it does require centralspindlin. In *C. elegans* embryos, disruption of the spindle midzone by SPD-1 (the *C. elegans* ortholog of PRC1) depletion does not block furrowing (Verbrugghe and White, 2004; Lee et al., 2015). However, these earlier experiments were performed in the presence of NOP-1, a nematode-specific ECT-2 activator (Tse et al., 2011). Therefore, we tested whether RhoA activation and furrowing can occur in embryos deficient in both SPD-1 and NOP-1. All such embryos initiated furrow formation, though ~50% of embryos did not complete furrow ingression. Furrow ingression was slower in the embryos depleted of SPD-1, but furrow initiation occurred more quickly following anaphase onset (Fig. 4F). The more rapid initiation of furrowing may be a consequence of the more rapid and extensive spindle elongation (Fig. 4G). In some embryos, RACGAP1 (CYK-4) was readily detected on the membrane during furrowing (Fig. 4H). These results indicate conservation of a spindle midzone-independent, and centralspindlin-dependent pathway for RhoA activation.

**Figure 2:**
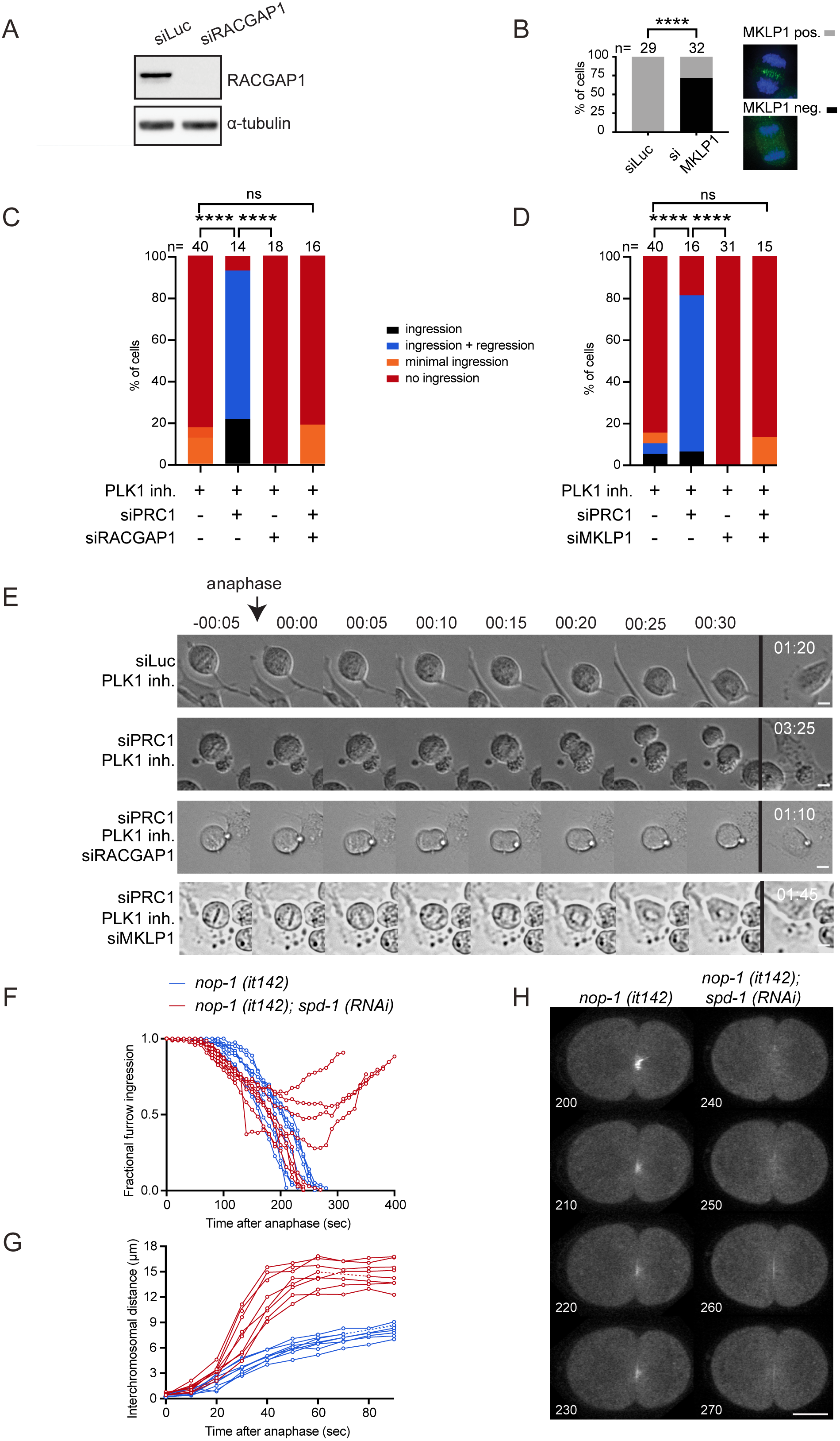
PLK1-independent furrow ingression requires centralspindlin. (A) Western blot of samples derived from HeLa cells transfected with either siLuc or siRACGAP1. The Western blot was probed with an anti-RACGAP1 antibody. α-Tubulin is shown as loading control. (B) HeLa cells were transfected with siLuc or siMKLP1 and processed for IF (right panel). The percentage of cells with detectable MKLP1 in anaphase was scored and used as a measure for knock-down efficiency. The number of cells that were scored per condition is indicated (n). One representative experiment out of 2 is shown. **** = p < 0.0001; Chi squared test for comparison of the indicated conditions. (C, D) HeLa cells transfected with the indicated siRNAs were imaged live. PLK1 was inhibited by addition of BI2536 (100 nM) at anaphase onset. The number of cells showing complete furrow regression, full furrow regression followed by regression, visible but minimal furrow ingression or no furrow ingression was scored. The number of cells analyzed per condition is indicated (n). One representative experiment out of 2 is shown. **** = p < 0.0001; Chi squared test for comparison of the indicated conditions, ns = not significant. (E) Representative DIC stills of HeLa cells transfected with the indicated siRNAs plus or minus addition of BI2536 (100 nM) prior to anaphase onset. Scale bar indicates 10 μm. (F,G) Quantification of the rate of furrow ingression (F) and chromosome separation (G) in *nop-1(it142)* embryos treated with *spd-1* or vector RNAi. Dotted lines reflect missing datapoints. Embryos expressed mCherry::Histone and mCherry::PH as well as CYK-4::GFP. (H) Stills of embryos imaged 200, 210, 220 and 230 sec. (vector RNAi), or 240, 250, 260 and 270 sec. (spd-1 RNAi) after anaphase onset. CYK-4::GFP is shown. Scale bar = 10 μm.

### Aurora B activity at the equatorial cortex is required for spindle midzone-independent furrow ingression

By depleting PRC1, we uncovered a PLK1-independent, but centralspindlin-dependent pathway for localized RhoA activation and furrow ingression. We hypothesized that another kinase might regulate centralspindlin-dependent RhoA activation and considered Aurora B kinase a likely candidate. Aurora B is located at the equatorial cortex in anaphase, and this localization becomes even more apparent after PRC1 depletion (Fig. 3A) (Earnshaw and Cooke, 1991; Murata-Hori and Wang, 2002; Mollinari et al., 2005). In both wild-type and PRC1-deficient cells, MKLP2 is required to translocate the kinase from the chromosomes to the equatorial cortex (Suppl. Fig. 2A,B) (Gruneberg et al., 2004; Hümmer and Mayer, 2009; Kitagawa et al., 2013). Moreover, we confirmed that this cortical localization of Aurora B was largely dependent on non-spindle midzone microtubules, because a concentration of nocodazole that did not depolymerize the stable spindle midzone microtubules, strongly reduced the localization of Aurora B at the equatorial cortex (Fig. 3B) (Théry et al., 2005; Murata-Hori and Wang, 2002; O’Connell and Wang, 2000). We next assessed the catalytic activity of this cortical pool of Aurora B. We fused a FRET-based Aurora B phosphorylation sensor to the c-terminal (aa 243-505) Tubby domain of mouse Tubby protein (Szentpetery et al., 2009; Santagata et al., 2001; Fuller et al., 2008), which binds phosphatidylinositol 4,5-biphosphate (PtdIns (4,5)P2) (Fig. 3C). The Tubby-FRET sensor was detected at the cell membrane before and during anaphase (Fig. 3C). Although chromosome-bound Aurora B is active during (pro)metaphase (Fuller et al., 2008), we did not detect phosphorylation of the membrane-localized sensor in early mitosis (Suppl. Fig. 2C). Phosphorylation of the membrane-localized sensor becomes detectable at anaphase onset, prior to visible membrane ingression, and was most prominent at the equatorial cortex (Fig. 3D-G). Depletion of MKLP2 or inhibition of Aurora B with ZM447439 before anaphase onset, prevented the generation of the signal from the FRET probe (Suppl. Fig. 2).

**Figure 3:**
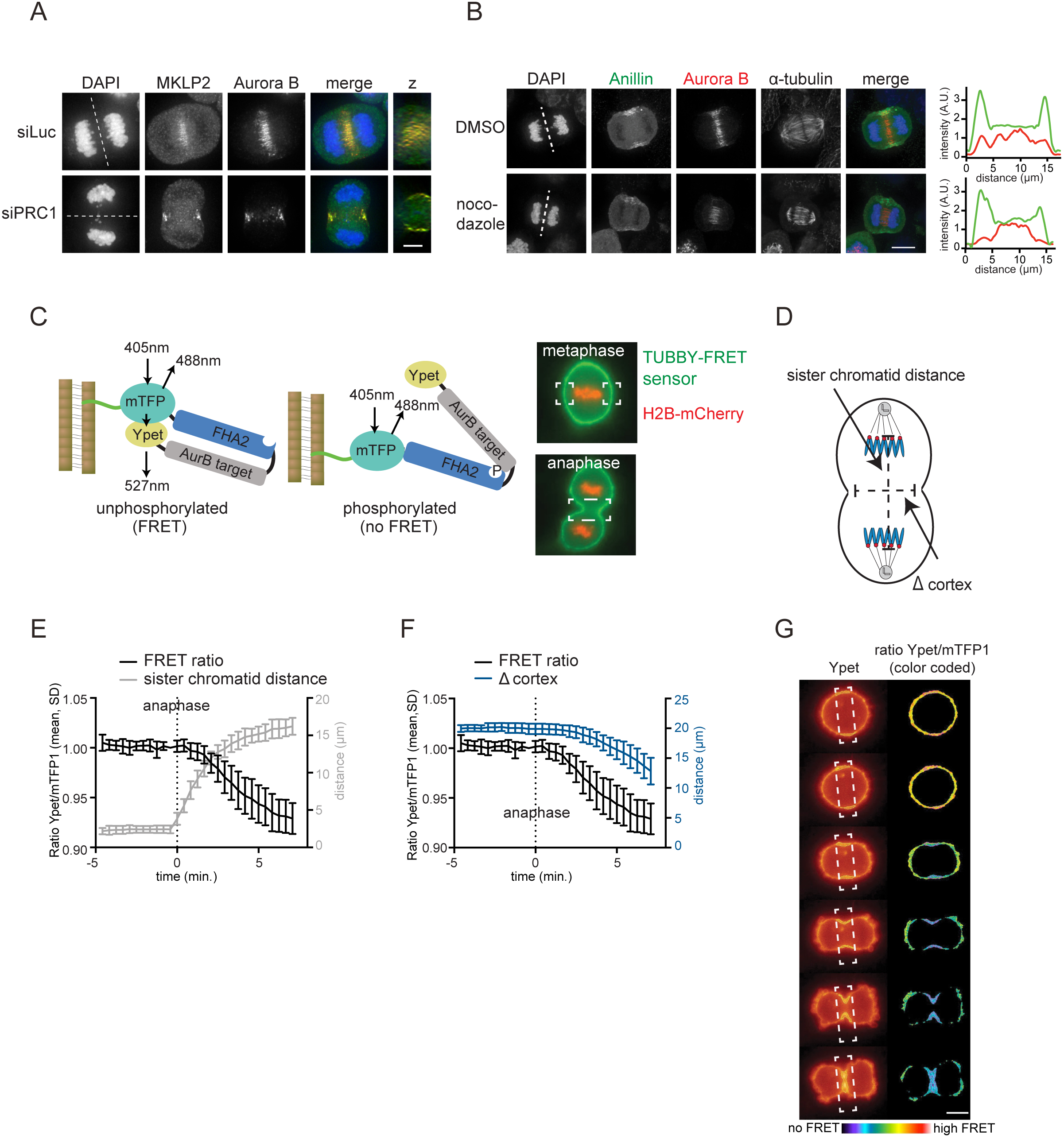
Aurora B localization and activity at the equatorial cortex. (A) IF for MKLP2 and Aurora B in HeLa cells transfected with siLuc or siPRC1. Dotted line indicates z-plane for z-axis view. Scale bar indicates 5 μm. (B) IF for Anillin and Aurora B in anaphase cells treated with 83 nM nocodazole to depolymerize microtubules that are not part of the spindle midzone. Scale bar indicates 5 μm. Dotted line indicates z-plane for line plot of Anillin (green) and Aurora B (red) intensity (far right). (C) Left: Scheme of the FRET-based Aurora B biosensor fused to Tubby protein (green line). Right: HeLa Flp-In T-Rex cells stably expressing Tubby-Aurora B FRET sensor (green) and H2B-mCherry (red). White boxes indicate the areas where FRET was measured. (D) Measurement of distance between the separating sister chromatids and the width of the ingressing furrow (*∆* cortex). (E, F) HeLa cells stably expressing the Tubby-Aurora B FRET sensor and H2B-mCherry were synchronized in G2 by treatment with the CDK1 inhibitor RO3306 and imaged live after release from the G2 block. The emission ratio at the equatorial cortex was calculated for each timepoint (interval = 25 sec., mean ± SD of 10 cells) and plotted with the distance between the separating sister chromatids (E) or with the width of the ingressing furrow (F). (G) Color-coded images of the Ypet/mTFP1 emission ratios. Scale bar = 10 μm.

Thus, an active pool of Aurora B kinase resides at the equatorial cortex at anaphase onset. To test the function of this pool of Aurora B, we inhibited Aurora B in both PRC1-proficient and PRC1-deficient cells. Inhibition of Aurora B activity in the presence of PRC1 did not prevent RhoA localization and Anillin accumulation at the equatorial cortex, and did not block cleavage furrow ingression (Fig. 4A-D). In line with previous studies, only the later stages of cytokinesis were impaired after Aurora B inhibition, as cleavage furrows eventually regressed (Fig. 4A,B) (Guse et al., 2005; Steigemann and Gerlich, 2009). However, in the absence of PRC1, cleavage furrow ingression was fully dependent on Aurora B activity (Fig. 4A-D). In addition, knock-down of MKLP2, which prevents the accumulation of Aurora B at the equatorial cortex (Suppl. Fig. 2A), also impaired furrow ingression in PRC1-depleted cells (Suppl. Fig. 2B). This suggests a pivotal role for localized Aurora B in stimulating RhoA activity at the equatorial cortex when the spindle midzone is disrupted (Fig. 4A-D). Of note, knockdown of PRC1 did not enhance Aurora B activity at the equatorial cortex (Suppl. Fig. 2F). This suggests that in the absence of PRC1, the spindle midzone-associated pool of Aurora B does not relocalize to the equatorial cortex to result in increased kinase activity. Thus, Aurora B activity is not critical for cytokinesis initiation in PRC1-proficient cells, but becomes essential for cleavage furrow ingression, after PRC1 depletion, when the spindle midzone is disrupted.

**Figure 4:**
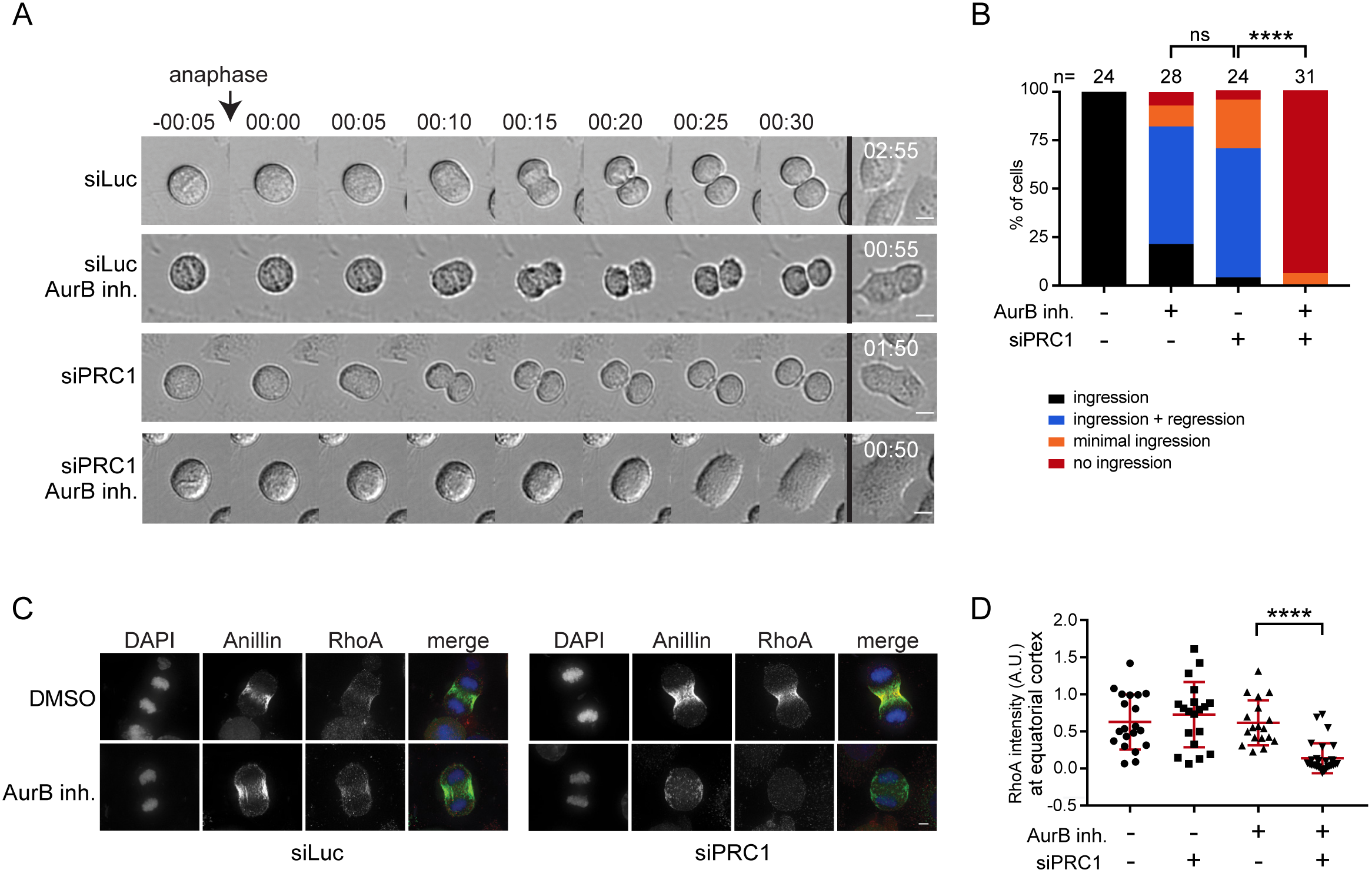
Aurora B activity at the equatorial cortex is required for spindle midzone-independent furrow ingression. (A) Representative DIC stills of a live cell imaging experiment with HeLa cells transfected with the indicated siRNAs and treated plus/minus the Aurora B inhibitor ZM447439 (2 μM) prior to anaphase onset. Scale bar indicates 10 μm. (B) Cells were treated and imaged as in (A), and the percentage of cells showing either stable furrow ingression, furrow ingression followed by furrow regression, minimal furrow ingression, no furrow ingression was scored. The number of cells analyzed per condition is indicated (n). One representative experiment out of 2 is shown. **** = p < 0.0001; Chi squared test for comparison of the indicated conditions, ns = not significant. (C) IF for Anillin and RhoA. DNA was visualized using DAPI. Scale bar = 5 μm. (D) Quantification of the fluorescence intensity levels of RhoA at the equatorial cortex in anaphase. Each dot represents an individual cell. Error bars depict the standard deviation (SD). **** = p < 0.0001; Student’s t-test for comparison of the indicated conditions.

### Aurora B-dependent centralspindlin oligomerization drives spindle midzone-independent furrow ingression

MKLP1 is an established substrate of Aurora B (Guse et al., 2005), and phosphorylation of S708 in MKLP1 by Aurora B disrupts the binding of 14-3-3 proteins to MKLP1 (Douglas et al., 2010). Dislodging of 14-3-3 from MKLP1 promotes centralspindlin oligomerization, which supports spindle midzone formation in human cells (Hutterer et al., 2009), and cortical contractility in *C. elegans* embryos (Basant et al., 2015) (Fig 5A). To test whether Aurora B-mediated centralspindlin oligomerization was responsible for spindle midzone-independent cytokinesis initiation in human cells, we generated a cell line stably expressing a phosphomimetic MKLP1-S708E variant (Fig. 5A-C). In otherwise wild-type cells, MKLP1 knockdown causes furrow regression after initial ingression, similar to what has been reported by others (Yüce et al., 2005;Nishimura, 2006; Zhao and Fang, 2005; Kamijo et al., 2006; Nguyen et al., 2014) (Fig. 5C). Expression of siRNA-resistant MKLP1-wt or MKLP1-S708E prevented furrow regression in the vast majority of the cells (Fig. 5B,C), confirming the functionality of the siRNA-resistant-GFP tagged MKLP1 proteins. Importantly, MKLP1-S708E rescued the initiation of cleavage furrow ingression in a substantial percentage of cells double depleted of PRC1 and MKLP1 and treated with the inhibitor of Aurora B (Fig. 5C). Collectively, this suggests that in PRC1 knockdown cells, Aurora B induces local clustering of centralspindlin that contributes to RhoA activation and implies that at least some centralspindlin resides at the equatorial cortex. To test the latter, we performed IF for MKLP1 and RACGAP1 in PRC1-depleted cells. Indeed, after PRC1 knock-down, MKLP1 and RACGAP1 no longer localized to microtubules in the midzone region (Fig. 5D) (Mollinari et al., 2005). However, we failed to detect MKLP1 at the equatorial cortex and only infrequently detected RACGAP1 at this site (Fig. 5D). Interestingly, in *C. elegans* embryos, the homolog of RACGAP1 (CYK4) can be detected on the equatorial plasma membrane as the furrow ingresses (Fig. 5E and Suppl. Movie 1). Depletion of Aurora B (AIR-2) in these cells results in a measurable loss of membrane accumulation of centralspindlin (Fig. 5E, F and Suppl. Movie 2). Based on these collective findings, we propose that in human cells a very small (below detection by IF), or a highly dynamic pool of centralspindlin at the equatorial cortex is sufficient to drive initiation of cytokinesis.

**Figure 5:**
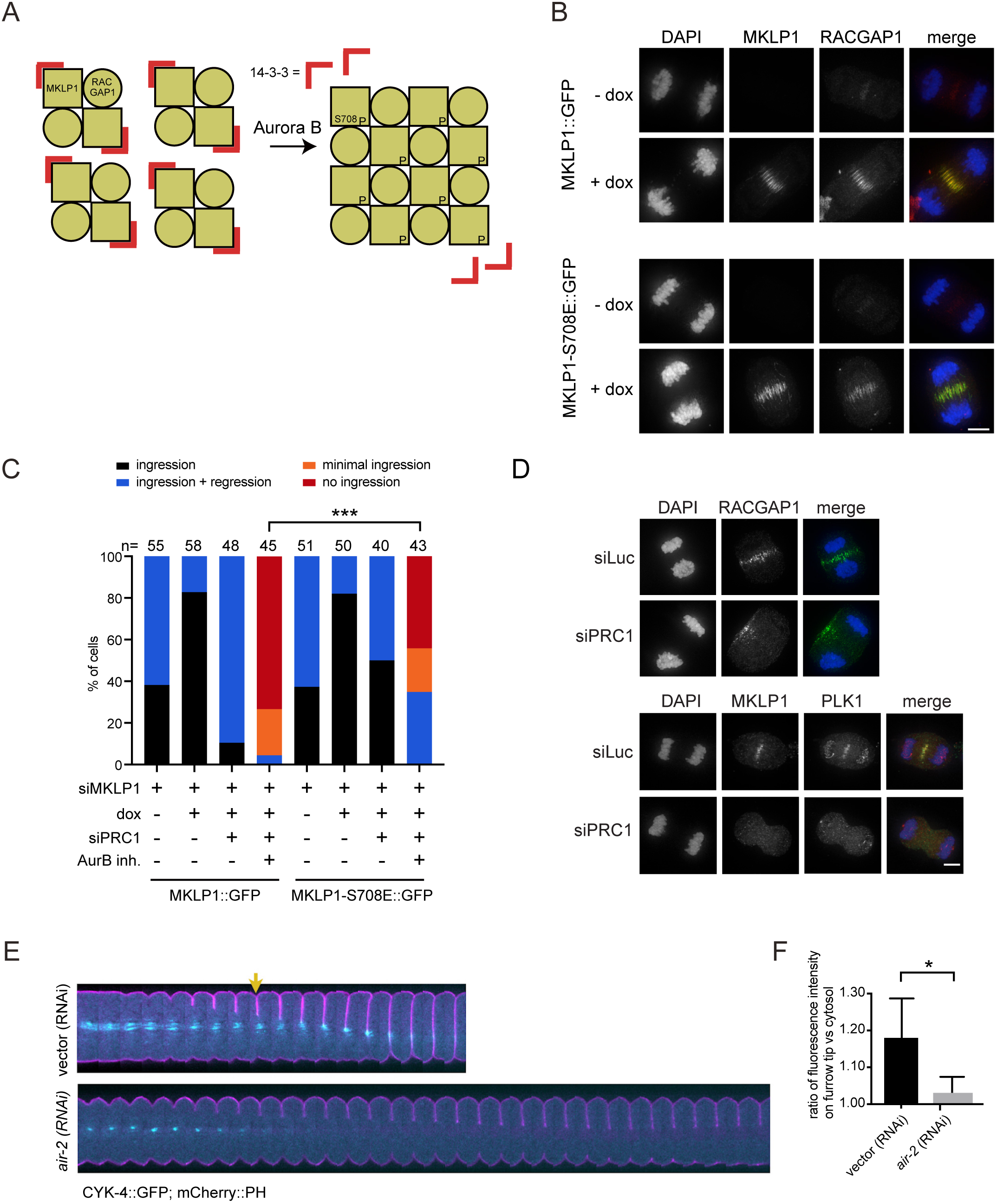
Aurora B-dependent centralspindlin oligomerization drives spindle midzone-independent furrow ingression. (A) Scheme explaining how oligomerization of centralspindlin is induced by Aurora B dependent phosphorylation of MKLP1. (B) HeLa cell lines with stable doxycycline (dox)-inducible expression of the indicated GFP-tagged (siRNA-insensitive) MKLP1 proteins were transfected with an siRNA for MKLP1 and processed for IF to visualize GFP-tagged MKLP1 and endogenous RACGAP1. For MKLP1::GFP expressing cells, 49/50 cells, and for the MKLP1-S708E::GFP expressing cells 47/51 cells showed GFP midzone localization. DNA was visualized using DAPI. Scale bar = 5 μm. (C) HeLa cell lines with stable inducible expression of MKLP1::GFP and MKLP1-S710E::GFP were transfected with the indicated siRNAs, treated with/without ZM447439 before anaphase, and imaged live. The number of cells showing complete furrow regression, full furrow regression followed by regression, visible but minimal furrow ingression or no furrow ingression was scored. The number of cells analyzed per condition is indicated. One representative experiment out of 2 is shown. *** = p < 0.001; Chi-squared test for comparison of the indicated conditions. (D) IF for RACGAP1, MKLP1 and PLK1 in cells transfected with siLuc or siPRC1. DNA was visualized using DAPI. Scale bar = 5 μm. (E) *C. elegans* embryos expressing a mCherry::PH membrane marker (pink) with a CYK-4::GFP transgene (cyan) were depleted of endogenous Aurora B (AIR-2) by RNAi. These embryos were filmed starting at metaphase in the first division cycle. Shown are montages of the equatorial region as the cell divides (n≥5). The arrow indicates the appearance of cortical CYK-4::GFP under wild-type conditions. Scale bar indicates 10 μm. (F) Embryos expressing CYK-4::GFP were scored for the extent of recruitment of the GFP marker to the ingressing furrow tip during anaphase, relative to a cytosolic background (see methods). Error bars represent ±SD, * = p < 0.05, Student’s t-test, n≥5.

### PLK1 suppresses PRC1 to allow anaphase spindle elongation and to limit sequestration of centralspindlin on the spindle midzone

Our findings suggest that RhoA activation at the equatorial cortex can occur by both PLK1-dependent and -independent cues, and that PLK1-independent RhoA activation requires Aurora B. If these cytokinesis initiation-routes act strictly in parallel, inhibition of one pathway would still allow the other pathway to operate. This is indeed observed after Aurora B inhibition, but not after PLK1 inhibition. Why does furrow ingression fail when PLK1 is inhibited in PRC1-proficient cells, when, according to our model, Aurora B should suffice to activate RhoA? One explanation could be that PLK1 inhibits cortical Aurora B activity when PRC1 is present. To test this possibility, we used the membrane-localized FRET-based Aurora B biosensor and found that Aurora B was equally active at the equatorial cortex in PLK1 inhibited cells as in cells with active PLK1 in which the furrow ingressed (Fig. 6A). We then asked whether PLK1 inhibition affects availability of Aurora B substrates at the cortex. PLK1 activity has been shown to suppress PRC1-dependent microtubule bundling and centralspindlin recruitment to the mitotic spindle in metaphase (Hu et al., 2012b). This prompted us to investigate whether PLK1 also limits PRC1 activity at the spindle midzone in anaphase. Indeed, spindle midzone microtubules are more bundled in PLK1-inhibited anaphase cells (Fig. 6B, E). This correlated with reduced anaphase spindle elongation after PLK1 inhibition, measured by a decrease in distance between the opposite cell poles, 5 min. before cell spreading (Fig. 6C) (Brennan et al., 2007). Midzone microtubule hyperbundling was diminished, and spindle elongation was restored after PRC1 depletion, suggesting that excessive crosslinking of midzone microtubules by PRC1 most likely interferes with anti-parallel microtubule sliding and that PLK1 is required to release this rigid state (Brennan et al., 2007; Lee et al., 2015). Moreover, quantification of the levels of MKLP1 and RACGAP1 at the spindle midzone revealed that the levels of these proteins increased when PLK1 is inhibited (Fig. 6D,E). Based on this, we conclude that PLK1 limits PRC1-mediated microtubule bunding in anaphase. We propose that when PLK1 is inhibited, PRC1 promotes excessive recruitment of centralspindlin, sequestering it away from the equatorial cortex. PRC1 depletion in PLK1-inhibited cells, releases centralspindlin from the spindle midzone allowing it to become activated by Aurora B at the equatorial cortex (Fig. 6F).

**Figure 6:**
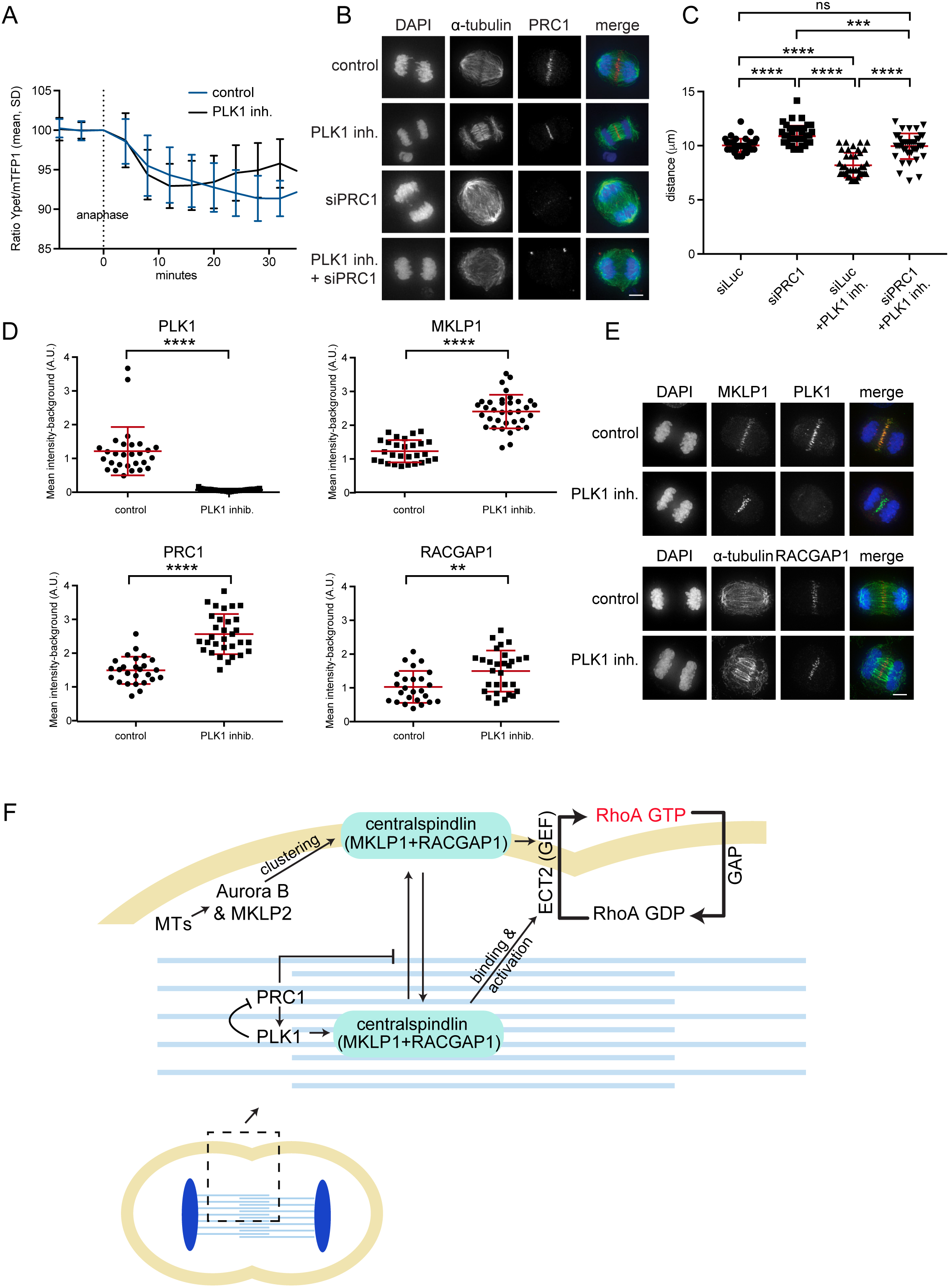
PLK1 suppresses PRC1 to allow dynamic exchange of centralspindlin between the spindle midzone and cell cortex. (A) HeLa cells stably expressing the Tubby-Aurora B FRET sensor and H2B-mCherry were synchronized in G2 using RO3306 and imaged live after release from the Cdk1 inhibitor. 35 minutes after release BI2536 (100 nM) was added. The emission ratio at the equatorial cortex was calculated for each time point (interval = 5 min). Mean ± SD of 11 cells is shown. (B) IF for α-tubulin and PRC1 of HeLa cells in anaphase transfected with the indicated siRNAs and treated plus/minus BI2536. Scale bar indicates 5 μm. (C) Cell elongation assessed one frame before cell spreading and measured as the cortex-cortex distance (based on DIC images) of opposite cell poles, perpendicular to the initial metaphase plate. Each dot represents and individual cell. One representative experiment out of 2 is shown. Error bars depict standard deviations. *** = p < 0.001; **** = p < 0.0001; Student’s t-test for comparison of the indicated conditions ns = not significant. (D) Quantification of the fluorescence intensity levels of PLK1, MKLP1, PRC1 and RACGAP1 at the spindle midzone in anaphase. Each dot represents an individual cell. One representative experiment out of 2 is shown. Error bars depict standard deviations. ** = p < 0.01; *** = p < 0.001, **** p < 0.0001 (Student’s t-test). (E) Representative images of C. Scale bar indicates 5 μm. (F) Model explaining how PLK1 and Aurora B activate centralspindlin and RhoA at the equatorial cortex. The phosphorylation of RACGAP1 by PLK1 promotes centralspindlin binding and activation of ECT-2, while Aurora B promotes ECT-2 activation via oligomerization of centralspindlin after phosphorylation of MKLP1. PLK1 also exerts an inhibitory effect on PRC1 which promotes the release of a fraction of centralspindlin from the spindle midzone. MTs = microtubules.

## Discussion

In this study we uncovered a PLK1-independent, Aurora B-dependent route to RhoA activation and cytokinesis initiation in human cells through knock-down of the microtubule bundling protein PRC1. This alternative PLK1-independent route to cytokinesis initiation has been missed, due to an unrecognized inhibitory effect of PLK1 on PRC1 in anaphase. We demonstrate that PLK1 constrains PRC1 in anaphase which serves two purposes; it limits PRC1’s microtubule bundling activity allowing spindle elongation, and second it promotes the release of a (small) pool of centralspindlin from the spindle midzone. This most likely allows centralspindlin to function both as a regulator of spindle midzone formation and as an activator of ECT-2 and RhoA at the equatorial cortex. How PLK1 constrains PRC1 remains to be fully determined. PRC1 is a direct substrate of PLK1, and its phosphorylation by PLK1 is needed to bind PLK1 and to localize the kinase on the spindle midzone (Neef et al., 2007; Hu et al., 2012a). This makes it inherently difficult to discriminate how PLK1 affects PRC1 function in anaphase, because mutation of the PLK1 phosphosites in PRC1 impairs PLK1 recruitment to the spindle midzone (Neef et al., 2007). PRC1 may directly sequester centralspindlin on the spindle midzone after PLK1 inhibition, as RACGAP1 can interact with PRC1 (Lee et al., 2015; Ban et al., 2004), or it may do so indirectly by creating hyper-bundled microtubules in the midzone. Distinguishing these possibilities through structure-function analysis of PRC1 is also challenging because its overexpression results in precocious spindle binding during metaphase (Mollinari et al., 2005; Hu et al., 2012b and our own unpublished observations).

An important implication from our work is that ECT-2 and RhoA activation can occur in the absence of PLK1 activity. PLK1 was shown to activate ECT-2 through phosphorylation of RACGAP1 which promotes its binding to the auto-inhibitory N-terminus of ECT-2. This relieves the intramolecular inhibition on the ECT-2 GEF domain (Wolfe et al., 2009; Burkard et al., 2009; Zhang and Glotzer, 2015). The N-terminus of RACGAP1 harbors 4 evolutionary conserved PLK1 phosphorylation sites and mutating these sites to alanine (RACGAP1-4A) attenuates complex formation between centralspindlin and ECT-2, fails to activate RhoA, and leads to loss of ECT-2 from the spindle midzone (Wolfe et al., 2009; Burkard et al., 2009). These results point to a crucial role for PLK1 in activating ECT-2 and RhoA. How can ECT-2 and RhoA become activated in PRC1-depleted cells when PLK1 is inhibited? Our results are complementary to recent work that demonstrates that mutations in the N-terminal BRCT domains of ECT-2 that strongly reduce its binding to RACGAP1 and its localization to the spindle midzone, do not prevent equatorial RhoA activity and cleavage furrow ingression and supported cytokinesis (Kotýnková et al., 2016). Importantly, knockdown of RACGAP1 still impaired furrow ingression in cells reconstituted with the RACGAP1 binding-deficient ECT-2 mutant (Kotýnková et al., 2016) as it does in PLK1-inhibited cells depleted of PRC1. Thus, RhoA activation during cytokinesis appears to be highly dependent on ECT-2 activation by centralspindlin. Notably, the ECT-2/RACGAP1 interaction is enhanced by, but does not require RACGAP1 phosphorylation by PLK1 (Somers and Saint, 2003; Yüce et al., 2005). RACGAP1 contains a C-terminal GAP domain that also interacts with ECT-2 which may contribute to formation and function of this complex (Zhang and Glotzer, 2015). This mode of ECT-2 and thereby RhoA activation might require, or be strongly promoted by, localized centralspindlin oligomerization at the equatorial cortex driven by Aurora B-dependent disengagement of 14-3-3 proteins from centralspindlin (Douglas et al., 2010; Basant et al., 2015).

This view is supported by optogenetic experiments with a C-terminally truncated ECT-2 that only activates RhoA at the equator (Kotýnková et al., 2016), presumably due to the presence of centralspindlin at this site. Moreover, active Aurora B is present at the equatorial cortex in PRC1-depleted cells and may provide a local environment for centralspindlin oligomerization. Indeed, in *C. elegans*, centralspindlin oligomerization obviates the requirement for Aurora B activity (Basant et al., 2015). In cultured cells, Aurora B activity is required for furrow ingression in PRC1-depleted cells and expression of MKLP1 S708E, mimicking Aurora B phosphorylation, partly restores ingression defects caused by Aurora B inhibition. This supports the idea that centralspindlin oligomerization can drive RhoA activation independent of PLK1. However, MKLP1 S708E, only rescues furrow ingression in ~30% of the PRC1-depleted and Aurora B inhibited cells. Most likely other Aurora B targets at the equatorial cortex, contribute furrow ingression in human cells (Goto et al., 2003; Neef et al., 2003).

One could argue that the main role of PLK1 in cytokinesis initiation is to limit PRC1 activity to make centralspindlin available for Aurora B-dependent activation at the equatorial cortex. However, in such a scenario, inhibition of Aurora B would always impair cytokinesis initiation and this is not the case: in PRC1-proficient cells, furrow ingression takes place when Aurora B is inhibited. This implies that PLK1-dependent RACGAP1 phosphorylation and activation of ECT-2 at the spindle midzone, and Aurora B-mediated oligomerization of centralspindlin at the equatorial cortex can in principle function as two separate pathways to centralspindlin and RhoA activation, and cytokinesis initiation. We propose that in wild-type cells, the “PLK1-brake” on PRC1 will also support the release of PLK1-phosphorylated centralspindlin bound to ECT-2, from the spindle midzone, allowing it to reach and activate RhoA at the equatorial cortex. Together, the PLK1- and Aurora B-dependent pathways to centralspindlin and RhoA activation may confer robustness to and proper timing of the processs of cleavage furrow ingression in mammalian cells.

In cells expressing the ECT-2 binding-deficient RACGAP-4A mutant, PLK1 is active and expected to act on PRC1 allowing the release of a fraction of centralspindlin from the spindle midzone that can then become activated at the equatorial cortex. In other words, the Aurora B-dependent route to furrow ingression should be operational. However, cleavage furrow ingression does not take place in the RACGAP1-4A expressing cells (Wolfe et al., 2009). Interestingly, the RACGAP1-4A mutant appears more concentrated at the spindle midzone (Wolfe et al., 2009), similar to what is observed for endogenous RACGAP1 after PLK1 inhibition. This raises the question whether RACGAP1 phosphorylation by PLK1 might also promote the release of centralspindlin from the spindle midzone.

In conclusion, we provide evidence for the existence of two pathways resulting in centralspindlin and RhoA activation and cytokinesis initiation in human cells. One pathway depends on PLK1 and originates at the spindle midzone, and the other pathway depends on Aurora B activity at the equatorial cortex. We argue that this latter pathway has gone unnoticed due to an unrecognized inhibitory effect of PLK1 on PRC1 in anaphase. We propose that the PLK1-dependent “brake” on PRC1 is necessary to release a fraction of centralspindlin from the spindle midzone that can activate ECT-2 and thus RhoA at the equatorial cortex. The finding that these two routes to centralspindlin and RhoA activation could operate independent from each other, highlights the robustness and plasticity of centralspindlin-induced cleavage furrow formation.

## Acknowledgements

We thank Dr. D. Gerlich (IMBA, Vienna) for the plasmid containing the Aurora B FRET sensor. This work is financially supported by the Netherlands Organization for Scientific Research (NWO-Vici 91812610 to S.M.A.L), by NIH R01GM085087 (to M.G.), and is part of the Oncode Institute which is partly financed by the Dutch Cancer Society.

## Supplemental Legends and Figures

**Supplemental Figure 1:**
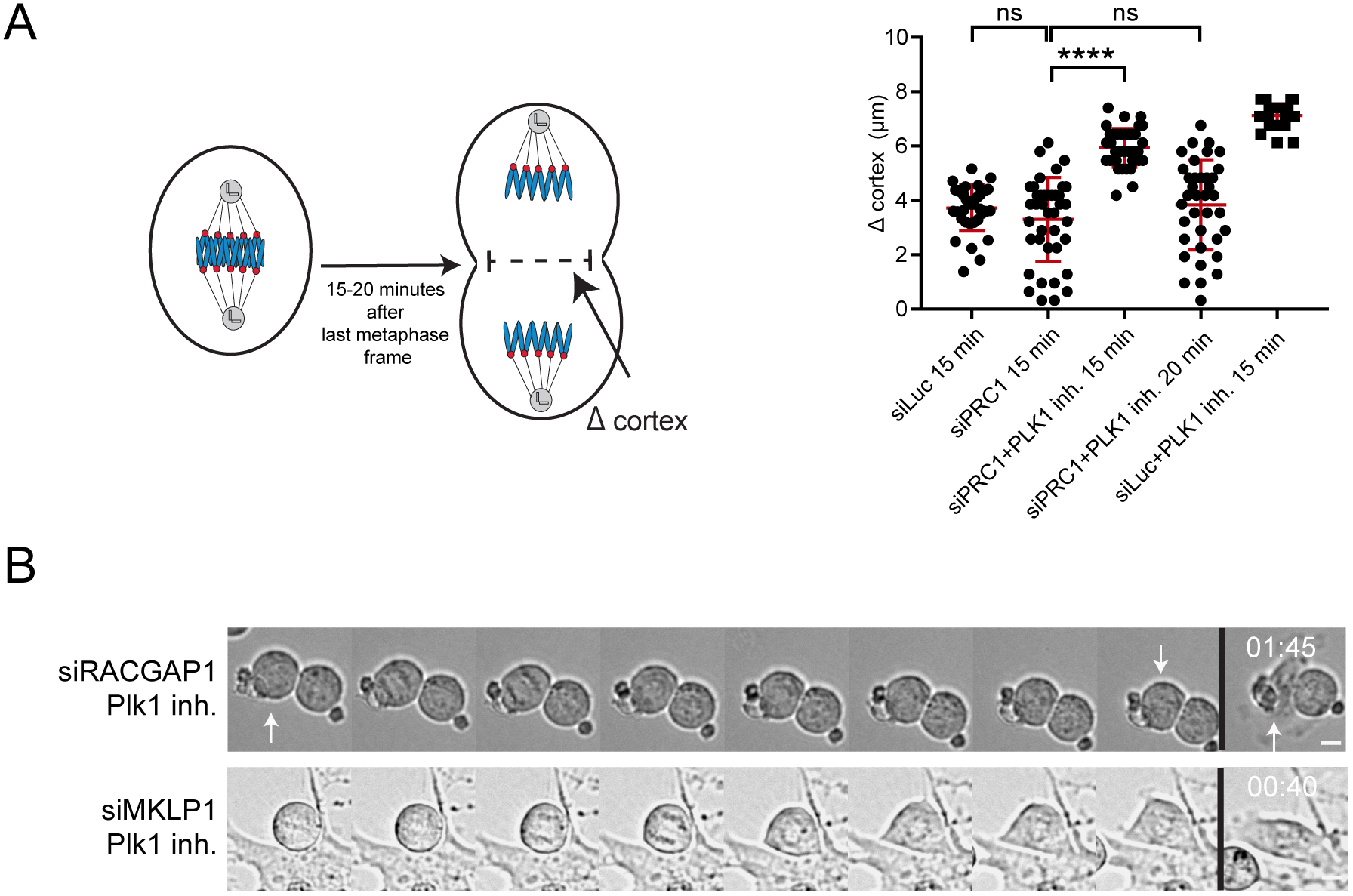
Furrow ingression in PLK1 inhibited HeLa cells. (A) Left: Scheme explaining how furrow ingression was measured 15-20 min. after the last frame in metaphase. Right: measurements of furrow ingression 15 min. after the last frame in metaphase. For the siPRC1 + PLK1 inh. condition also a 20 min. timepoint was measured. Each dot represents an individual cell. Error bars depict standard deviations. **** p < 0.0001; Student’s t-test for comparison of the indicated conditions, ns = not significant. (B) Representative DIC stills of a live cell imaging experiment of HeLa cells treated with indicated siRNAs plus or minus addition of BI2536 (100 nM) prior to anaphase onset. Scale bar = 10 μm.

**Supplemental Figure 2:**
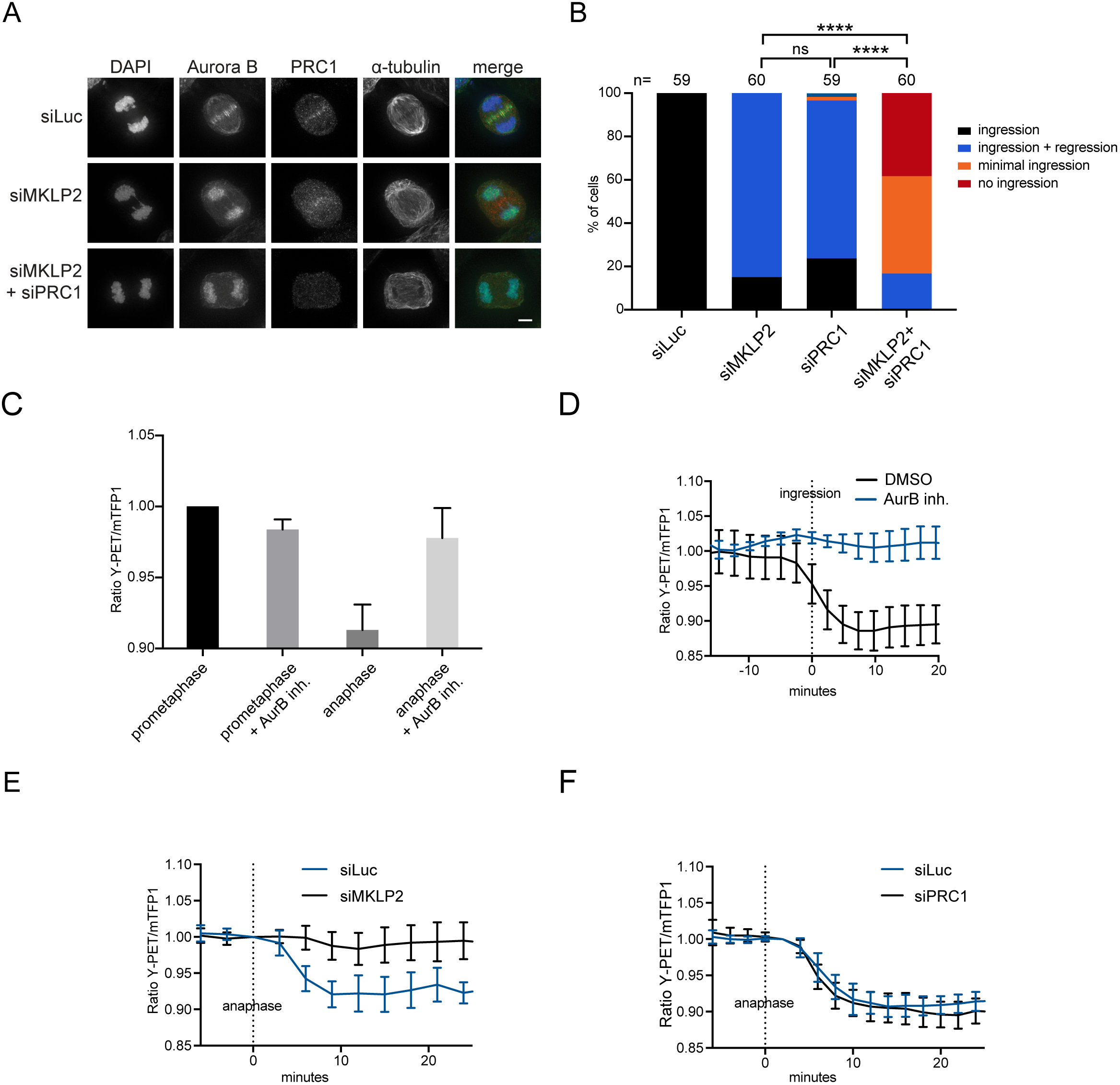
Aurora B localization and activity at the equatorial cortex. (A) IF for α-tubulin, PRC1 and Aurora B of HeLa cells in anaphase transfected with the indicated siRNAs. Scale bar = 5 μm. (B) HeLa cell lines were transfected with the indicated siRNAs and imaged live. The number of cells showing complete furrow regression, full furrow regression followed by regression, visible but minimal furrow ingression or no furrow ingression was scored. The number of cells analyzed is indicated (n). One representative experiment out of 2 is shown. **** = p < 0.0001; Chi-squared test for comparison of the indicated conditions, ns = not significant. (C,D) HeLa cells stably expressing the Tubby-Aurora B FRET sensor and H2B-mCherry were synchronized in G2 using RO3306 and imaged live after release from the Cdk1 inhibitor. Thirty-five minutes after RO3306 release, DMSO or the Aurora B inhibitor ZM447439 was added. (C) The emission ratio at the equatorial cortex was calculated in prometaphase (~17 min before furrow ingression) and in anaphase (~3 min after furrow ingression). Data were derived from experiment presented in D. (D) The emission ratio at the equatorial cortex was calculated for each time point (interval = 3 min). Mean ± SD of 11 (DMSO), or 10 (ZM447439) cells is shown. (E) HeLa cells stably expressing the Tubby-Aurora B FRET sensor and H2B-mCherry were treated with control (siLuc) or a MKLP2 depletion and were synchronized in G2 using RO3306 and imaged live after release from the Cdk1 inhibitor. The emission ratio at the equatorial cortex was calculated for each time point (interval = 3 min). Mean ± SD of 15 (siLuc) and 14 (siMKLP2) cells is shown. (F) HeLa cells stably expressing the Tubby-Aurora B FRET sensor and H2B-mCherry were transfected with siLuc or siPRC1, synchronized in G2 using RO3306, and imaged live after release from the CDK1 inhibitor. The emission ratio at the equatorial cortex was calculated for each time point (interval = 3 min). Mean ± SD of 6 (siLuc) and 9 (siPRC1) cells is shown.

Supplemental Movie 1: *C.elegans* embryos expressing mCherry::PH membrane marker and CYK4::GFP were filmed starting at metaphase in the first division cycle. Only CYK4::GFP is shown in the movie. Montages with mCherry::PH are shown in Figure 5E.

Supplemental Movie 2: *C.elegans* embryos expressing mCherry::PH membrane marker and CYK4::GFP were depleted of endogenous Aurora B (AIR-2) by RNAi and filmed starting at metaphase in the first division cycle. Only CYK4::GFP is shown in the movie. Montages with mCherry::PH are shown in Figure 5E.

